# A novel QTL conferring Fusarium crown rot resistance located on chromosome arm 6HL in barley

**DOI:** 10.1101/537605

**Authors:** Shang Gao, Zhi Zheng, Haiyan Hu, Haoran Shi, Jian Ma, Yaxi Liu, Yuming Wei, You-Liang Zheng, Meixue Zhou, Chunji Liu

## Abstract

Fusarium crown rot (FCR), caused primarily by *Fusarium pseudograminearum*, is a devastating disease for cereal production in semi-arid regions worldwide. To identify and characterize loci conferring FCR resistance, we assessed a landrace AWCS799 which is among the top lines identified from a screening. Genetic control of its resistance was investigated by generating and analysing two populations of recombinant inbred lines. One of the populations was used for QTL detection and the other for validation. A novel QTL, located on the long arm of chromosome 6H (designated as *Qcrs.caf-6H*), was consistently detected in each of the four trials conducted against the mapping population. The QTL explained up to 29.1% of the phenotypic variance and its effect was confirmed in the validation population. Significant interactions between this resistance locus and either plant height or heading date were not detected, further facilitating its manipulation in breeding programs.

**Key Message:** This study identified and validated a novel and large-effect QTL conferring Fusarium crown rot resistance on the long arm of chromosome 6HL in barley.

## Introduction

Fusarium crown rot (FCR) is a chronic disease caused by various *Fusarium* species in cereal crops. It is prevalent in arid and semi-arid cropping regions worldwide (Akinsanmi et al. 2004; Chakraborty et al. 2006). The disease has been known for causing significant yield loss in Australia (Murray and Brennan 2010; Murray and Brennan 2009) and the USA (Smiley et al. 2005b). FCR has also become a major issue in recent years for cereal production in China (Li et al. 2016; Xu et al. 2017; Zhou et al. 2014).

FCR pathogens are carried over in crop residues. They can survive for two seasons or longer in the field environments (Chakraborty et al. 2006; Smiley et al. 2005a) making the disease difficult to manage using practices such as crop rotation (Burgess 2014). As recognised several decades ago, growing resistant varieties has to be an integral component in effectively managing the disease (Purss 1966; Wildermuth and Purss 1971). Growing resistant varieties does not only reduce yield losses but could also reduce the loss of the following barley or other cereal crops by reducing the inoculum load. This is especially the case for barley as, compared with those of wheat, its plants accumulate much higher concentrations of *Fusarium* pathogens at every stage of CR infection (Liu et al. 2012a).

The availability of well-characterized genotypes with high levels of resistance would facilitate breeding varieties with enhanced resistance. To date, there are only three reported studies on identifying QTL conferring FCR resistance in barley. The first one was based on a study using an existing population developed for unrelated traits. The study detected a single locus on chromosome arm 3HL which interacts strongly with plant height (Li et al. 2009). Effects of plant height on FCR development were also detected based on near isogenic lines (Chen et al. 2014) and histological analyses (Bai and Liu 2015). The second study detected a single locus on chromosome arm 4HL (designated as *Qcrs.cpi-4H*) from a wild barley genotype (*Hordeum spontaneum* L.) originated from Iran (Chen et al. 2013a). The third study detected two loci from a landrace originated from Japan. They located on 1HL (*Qcrs.cpi-1H*) and 3HL (*Qcrs.cpi-3H*) (Chen et al. 2013b).

Gene pyramiding has shown to be effective in further improving FCR resistance in both barley (Chen et al. 2015) and wheat (Zheng et al. 2017). Understandably, the quality and quantity of available loci are critical in determining the effectiveness of gene pyramiding or breeding programs. With the aim of identifying additional loci with high levels of resistance to FCR, we conducted a QTL mapping study against a landrace originating from South Korea. It was identified as one of the most resistant genotypes from a systematic screening of more than 1,000 genotypes representing diverse geographical origins and different plant types (Liu et al. 2012b). As previous studies have repeatedly shown that both plant height (Bai and Liu 2015; Chen et al. 2013a; Li et al. 2009; Ma et al. 2010; Zheng et al. 2014) and heading date (Liu et al. 2012b) may interact with FCR severity, we investigated possible interactions between QTL detected from this study with these characteristics. Results obtained from these analyses are reported in this publication.

## Materials and Methods

### Plant materials

The genotype AWCS799 is the resistant source analysed in this study. Two populations of recombinant inbred lines (RILs) between ‘AWCS799’ and two cultivars were developed and used in this study. They are:

1. Fleet/AWCS799 consisting of 124 F8 RILs.
2. Franklin/AWCS799 consisting of 121 F8 RILs.

Both populations were produced in glasshouses at the Queensland Bioscience Precinct (QBP) in Brisbane, Australia. The first population was used for QTL mapping and the other for validating putative QTL identified from the mapping population.

### FCR inoculation and disease assessments

Results from previous studies show that FCR resistance is not pathogen species-specific and the same resistance locus can be detected by pathogen isolates belonging to different *Fusarium* species (Chen et al. 2013b; Ma et al. 2010). We therefore used a single *F. pseudograminearum* isolate in this study. The isolate (CS3096) was obtained from a wheat field in northern New South Wales, Australia and maintained in the CSIRO collection (Akinsanmi et al. 2004). Inoculum preparation, inoculations, and FCR assessments were as described by Li et al. (2008). Briefly, inoculum was prepared using plates of ½ strength potato dextrose agar. Inoculated plates were kept for 12 days at room temperature before the mycelium in the plates were scraped and discarded. The plates were then incubated for a further 7-12 days under a combination of cool white and black fluorescent lights with 12-h photoperiod. The spores were then harvested and the concentration of spore suspension was adjusted to 1×10^6^ spores/ml. The spore suspensions were stored in minus 20 freezer and Tween 20 was added (0.1%v/v) to the spore suspension prior to use.

Seeds were germinated in Petri dishes on three layers of filter paper saturated with water. Seedlings of 3-day-old were immersed in the spore suspension for 1 min and two seedlings were planted into a 3 cm square punnet (Rite Grow Kwik Pots, Garden City Plastics, Australia) containing sterilized University of California mix C (50% sand and 50 % peat v/v). The punnets were arranged in a randomized block design and placed in a controlled environment facility (CEF). Settings for the CEF were: 25/18 (±1) °C day/night temperature and 65/80% (±5)% day/night relative humidity, and a 14-h photoperiod with 500 μmol m^−2^S^−1^ photon flux density at the level of the plant canopy. To promote FCR development, water-stress was applied during plant growth. Inoculated seedlings were watered only when wilt symptoms appeared.

For the initial QTL mapping, four replicated trials were carried out against the mapping population (designated as FCR01 to FCR04, respectively). Three replicated trials were conducted on the validation population (designated as FCRV01, FCRV02, FCRV03, respectively). Fourteen seedlings were used for each of the replicates.

FCR severity was assessed 4 weeks after inoculation, using a 0 (no visible symptom) to 5 (whole plant severely to completely necrotic) scale as described by Li et al. (2008). A disease index (DI) was them calculated for each line following the formula of DI = (ΣnX/5 N) × 100, where X is the scale value of each plant, n is the number of plants in the category, and N is the total number of plants assessed for each line.

The graph for the frequency distribution of DI values and correlations between all phenotypic data were generated with R package ‘ggplot2’ (Wickham 2016) and ‘psych’ (Revelle 2017), respectively. The best linear unbiased prediction (BLUP) for the DI value was calculated using ‘lmer’ function in the R package ‘lme4’ (Bates et al. 2007). Student’s *t* test performed by Microsoft Office Excel 2016 was used to evaluate the difference in DI values between lines with or without the resistant alleles in the populations.

### Evaluation for plant height and heading date

To assess possible effects of plant height (PH) and heading date (HD) on FCR resistance, two trials were conducted using the Fleet/AWCS799 population at the CSIRO Research Station at Gatton, Queensland (27°34′S, 152°20′E). A randomized block design was used with three replicates for each of the field trials. For each replicate, 20 seeds for each of the RILF_8_ lines were sown in a single 1.5 metre row with a 25 cm row-spacing. PH was measured as the height from the soil surface to the tip of the spike (awns excluded). Six measurements were taken from the six tallest tillers in each row and the average of each line were used for statistical analyses. Heading date was recorded for each line on which about 50% of the spikes emerged from main tillers.

### Molecular marker analysis

Genotypes for the two parents and 94 RILs from the population Fleet/AWCS799 were generated by the Department of Economic Development, Jobs, Transport and Resources (DEDJTR), Victoria, Australia, according to a tGBS (an optimized approach for genotyping-by-sequencing) pipeline (Ott et al. 2017). SSR markers were then developed for putative QTL regions and used to genotype the whole mapping population.

PCR reactions for the amplification of the SSR markers were carried out in a total volume of 12 μl containing 25 ng genomic DNA, 0.2 μM of forward and reverse primer, 3 mM MgCL_2_, 0.2 mM dNTPs, and 0.5 U *Taq* DNA polymerase. During PCR, marker products were labelled with α-[^33^P]dCTP (3,000 ci/mmol). PCR reactions were run on a Gene Amp PCR system 2700 thermocycler (PE Applied Biosystems, Foster City, CA, USA) programmed with the cycling conditions: one cycle of 5 min at 94 °C, 35 cycles of 30 sec at 94 °C, 30 sec at 60 °C (annealing temperature for 6H-2) and 1 min at 72 °C, with a final extension step of 5 min at 72 °C. The amplified products were mixed with an equal volume of loading dye, denatured at 95 °C for 5 min, and 4 μl samples was run on a denaturing 5% polyacrylamide (20:1) gel at 110 W for 2 h. The gels were subsequently dried using a gel dryer for 30 min at 80 °C and exposed to Kodak X-Omat X-ray films for 2 days.

### Data analysis and QTL mapping

Statistical analyses were performed using the Microsoft Office Excel 2016 and R. For each of the trials, the following mixed-effect model was used: Y_*ij*_ = μ + r_*i*_ + g_*j*_ + w_*ij*_. Where: Y_*ij*_ =trait value on the jth genotype in the ith replication; μ =general mean; r_*i*_ = effect due to ith replication; g_*j*_ = effect due to the jth genotype; w_*ij*_ =error or genotype by replication interaction, where genotypes was treated as a fixed effect and that of replicate as random. The effects of replicate and genotype for both FCR resistance and PH were determined using ANOVA. Pearson correlation coefficient was estimated between the traits and trials.

MSTmap Online (Wu et al. 2008) was used to build linkage maps by chromosomes with the following parameters: Grouping LOD Criteria: single LG; Population type: RIL8; No mapping missing threshold: 10%; No mapping distance threshold: 15 cM; No mapping size threshold: 2; Try to detect genotyping errors: Yes; Genetic mapping function: Kosambi. The diagrams of linkage maps were generated with MapChart (Voorrips 2002). IciMapping 4.1 was used for QTL analysis with the “Biparental Population” (BIP) module (Meng et al. 2015). Interval mapping (IM) and Inclusive composite interval mapping (ICIM) was then employed to identify QTL. For each trial, a test of 1,000 permutations was performed to decide the LOD threshold corresponding to a genome-wide type I error less than 5% (P < 0.05). QTL were named according to the rules of International Rules of Genetic Nomenclature (http://wheat.pw.usda.gov/ggpages/wgc/98/Intro.htm).

## Result

### Characterization of FCR resistance in the mapping population of Fleet/AWCS799

FCR severity of the resistant genotype AWCS799 was significantly lower than the two susceptible commercial cultivars (Fig. 1). In the four trials conducted against the mapping population and its two parents, the DI value of AWCS799 was 32.6% lower than that of Fleet on average and transgressive segregation was detected in each of the trials (Table 1). DI values for all four trials and BLUP presented significantly positive correlation between each other (Table 2). The frequency distribution of DI value for FCR01 skewed towards better resistance. DI values for the other three trials and BLUP showed more normal distributions (Fig. S1).

**Fig. 1.**
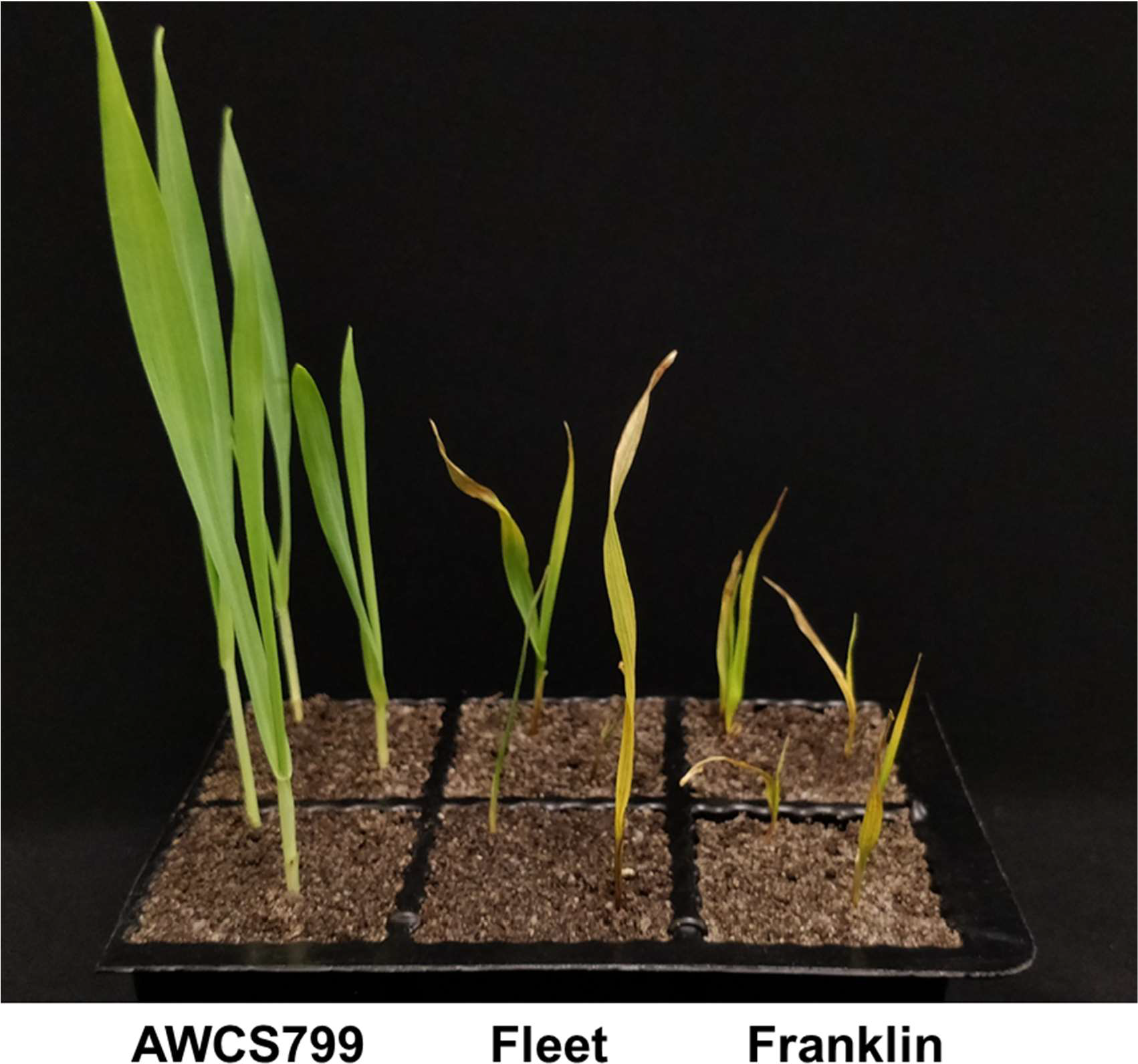
Difference in resistance to Fusarium crown rot infection between the resistant genotype AWCS799 and the two commercial cultivars (Fleet and Franklin) used as parents in this study

**Table 1.** Disease index of FCR severity in the population of Fleet/AWCS799

| Trial | Parent |  | Population |  |  |  |
| --- | --- | --- | --- | --- | --- | --- |
|  | Fleet | AWCS799 | Min | Max | Mean | SD |
| FCR01 | 55.0 | 38.4 | 25.5 | 67.2 | 41.7 | 10.3 |
| FCR02 | 55.4 | 35.8 | 28.6 | 76.2 | 50.1 | 9.4 |
| FCR03 | 58.6 | 43.4 | 28.6 | 73.0 | 47.1 | 9.4 |
| FCR04 | 66.1 | 40.2 | 31.4 | 69.1 | 50.8 | 8.8 |
| BLUP | 55.8 | 34.3 | 29.5 | 64.5 | 45.1 | 7.0 |

**Table 2.** Correlation coefficients between FCR severity, plant height and heading date in the Fleet/AWCS799 population

|  | FCR01 | FCR02 | FCR03 | FCR04 | BLUP | PH | HD |
| --- | --- | --- | --- | --- | --- | --- | --- |
| FCR01 | 1.00 |  |  |  |  |  |  |
| FCR02 | 0.23** | 1.00 |  |  |  |  |  |
| FCR03 | 0.88* | 0.29** | 1.00 |  |  |  |  |
| FCR04 | 0.33** | 0.25** | 0.36** | 1.00 |  |  |  |
| BLUP | 0.79** | 0.56* | 0.83** | 0.66** | 1.00 |  |  |
| PH | -0.11 | -0.10 | -0.06 | -0.10 | -0.13 | 1.00 |  |
| HD | -0.10 | -0.11 | 0.03 | 0.06 | -0.08 | 0.35** | 1.00 |

‘\*’ significant at $p < 0.05$ ; ‘\*\*’ significant at $p < 0.01$ ; ‘PH’ stands for plant height; ‘HD’ stands for heading date.

### Linkage maps constructed and synteny for marker locations in the genome assembly of Morex

Of the GBS markers mapped, 4,870 codominant markers detected polymorphism between Fleet and AWCS799. These markers fell into 740 clusters and markers within each of the clusters co-segregated. As co-segregating markers contain the same information when used for mapping, a single marker with the least missing values was selected from each of the clusters and used for linkage map construction.

The markers were grouped into seven linkage groups and they spanned a total of 1964.7 cM with an average distance of 2.3 cM between loci (Table S1, Fig. S2). As all of the markers generated had known physical positions, we aligned the linkage maps with the latest genome assembly of barley reference genotype Morex. This analysis found that, as expected, the genetic and physical maps were highly consistent for majority of the markers. However, there are a few exceptions and, interestingly, they were all located in the peri-centromeric regions (Fig. S3).

### Detection and validation of QTL for FCR resistance

Four putative QTL were detected from the mapping population. They were located on chromosomes 2H, 4H, 5H and 6H (Fig. 2), respectively. Only the one located on 6H was detected in each of the four trials conducted. The resistant allele of this QTL originated from AWCS799. We designated this QTL as *Qcrs.caf-6H*, where ‘Crs’ stands for ‘crown rot severity’ and ‘caf’ represents ‘CSIRO Agriculture and Food’. Qcrs.*caf-6H* was identified in all four trials and BLUP using both IM and ICIM analysis and it could explain up to 29.1% of the phenotypic variance (Fig. 3; Table 3). Based on ICIM analysis, *Qcrs.caf-6H* was narrowed to a 4.0 cM-interval flanked by markers 6H_478006625 and 6H_481998837 with 6H_497772849 (forward primer: 5’-GCATTAGTTGTCATAGTAGGTAGCA-3’; reverse primer: 5’-TTCAAGACCACGACCTTGGG-3’) as the most closely linked SSR marker.

**Fig. 2.**
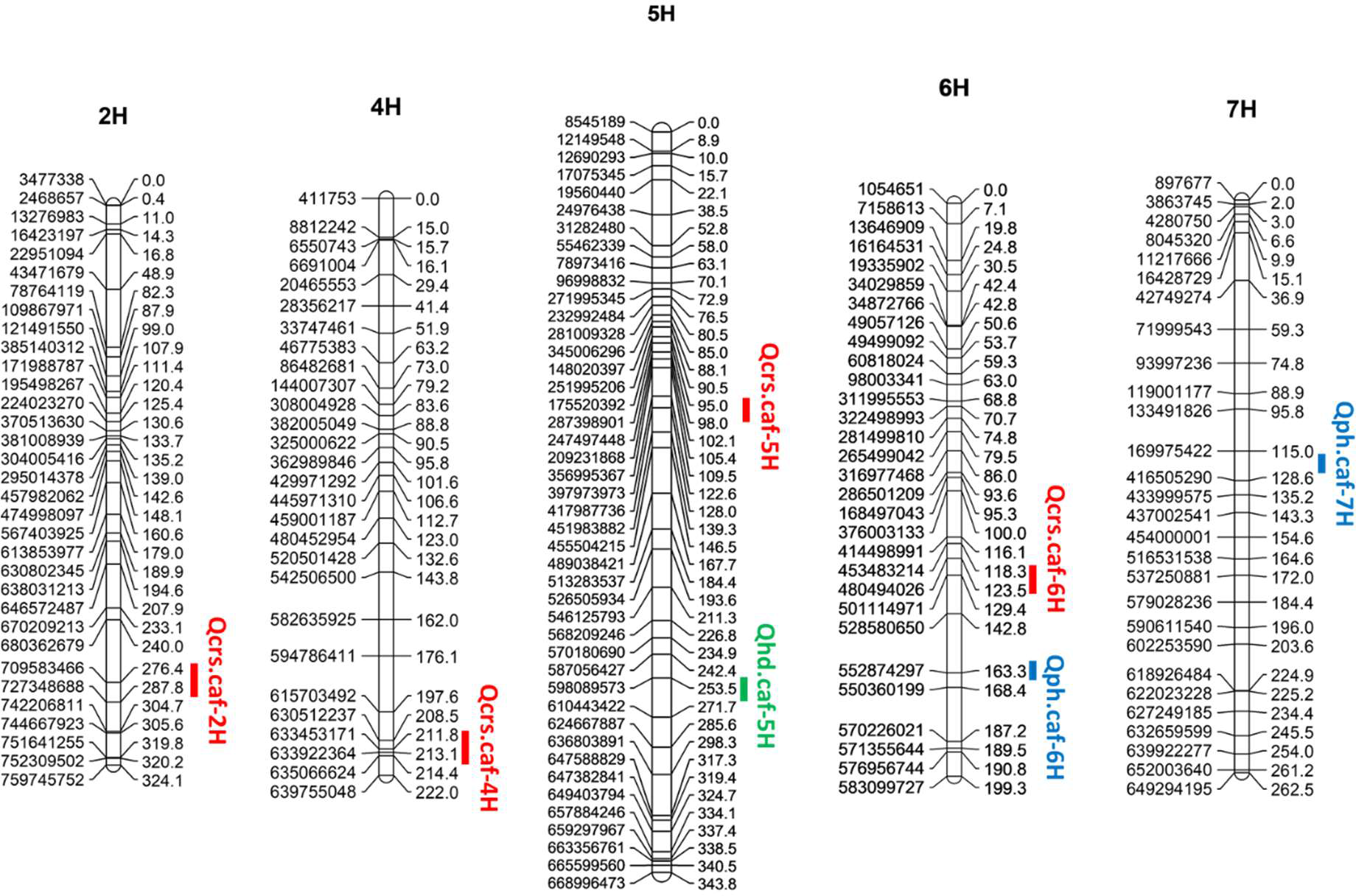
QTL conferring FCR resistance (red), plant height (blue) and heading date (green) detected from the population of Fleet/AWCS799. Physical position for each marker is shown to the left of the linkage map and distances in centiMorgan (cM) are shown to the right.

**Fig. 3.**
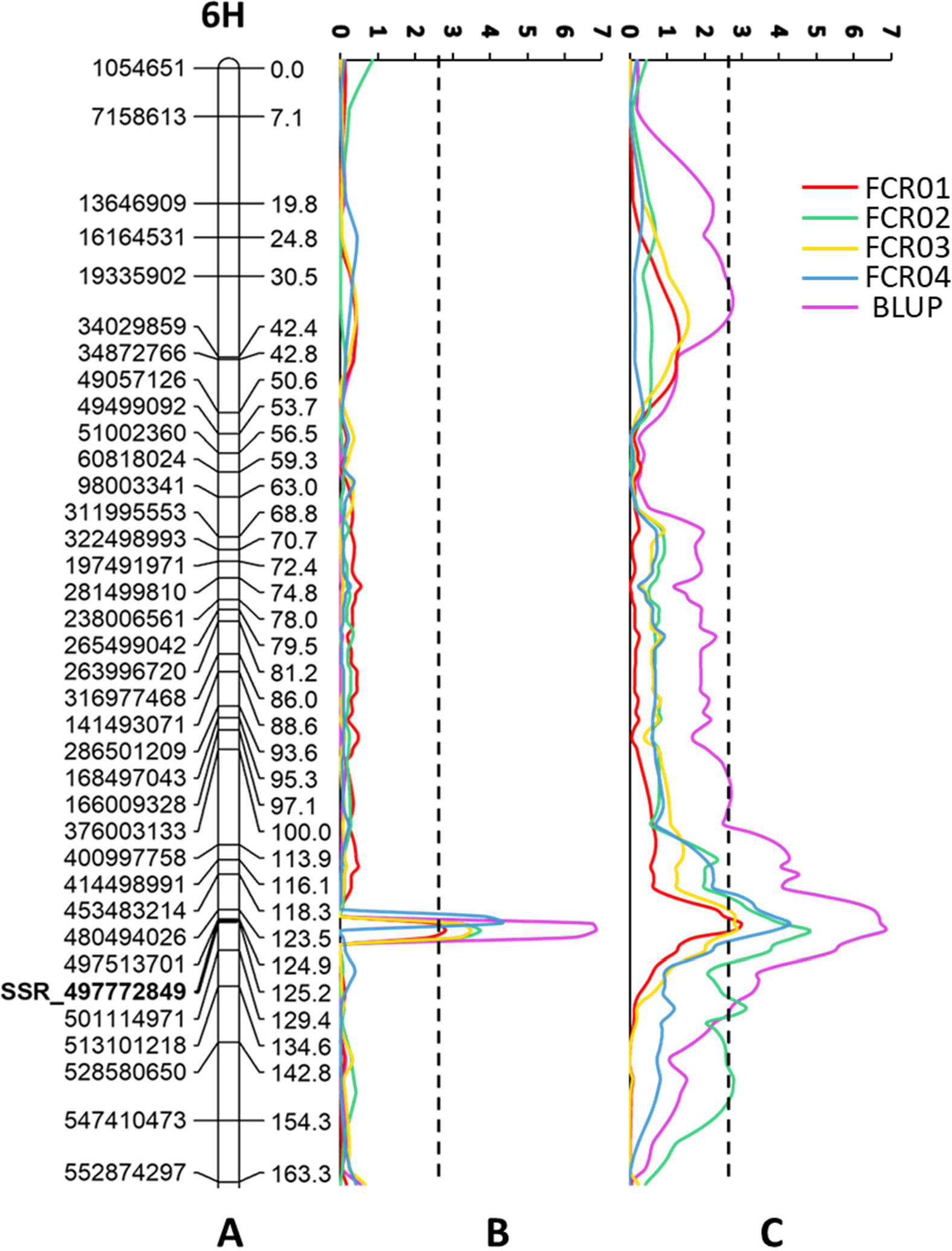
QTL conferring FCR resistance detected on the long arm of chromosome 6H in the population of Fleet/AWCS799. A: Physical position for each marker is shown to the left of the linkage map and distances in centiMorgan (cM) are shown to the right; B: LOD value graph for “ICIM” model; C: LOD value graph for “IM” model. The vertical dotted lines indicate the average significance threshold (LOD=2.7) based on a test of 1,000 permutations. The SSR marker used in validating the QTL is bolded.

**Table 3.** QTL for FCR severity, plant height and heading date identified in the population of Fleet/AWCS799

| QTL | Trials <sup>a</sup> | Analysis <sup>b</sup> | LOD | Interval | Flanking markers | PVE (%) <sup>c</sup> | Origin |
| --- | --- | --- | --- | --- | --- | --- | --- |
| *Qcrs.caf-6H* | FCR01 | ICIM | 2.8 | 122.5 - 125.5 | 6H_480494026 & 6H_481998837 | 13.1 | AWCS799 |
|  | FCR02 | ICIM | 3.8 | 122.5 - 125.5 | 6H_480494026 & 6H_481998837 | 16.5 | AWCS799 |
|  | FCR03 | ICIM | 3.5 | 122.5 - 125.5 | 6H_480494026 & 6H_481998837 | 15.7 | AWCS799 |
|  | FCR04 | ICIM | 4.3 | 121.5 - 123.5 | 6H_478006625 & 6H_480494026 | 21.6 | AWCS799 |
|  | BLUP | ICIM | 6.8 | 122.5 - 125.5 | 6H_480494026 & 6H_481998837 | 29.1 | AWCS799 |
|  | FCR01 | IM | 3.0 | 123.5 - 127.5 | 6H_480494026 & 6H_481998837 | 11.4 | AWCS799 |
|  | FCR02 | IM | 4.8 | 122.5 - 126.5 | 6H_480494026 & 6H_481998837 | 9.1 | AWCS799 |
|  | FCR03 | IM | 2.9 | 123.5 - 127.5 | 6H_480494026 & 6H_481998837 | 11.4 | AWCS799 |
|  | FCR04 | IM | 4.3 | 120.5 - 124.5 | 6H_478006625 & 6H_480494026 | 9.7 | AWCS799 |
|  | BLUP | IM | 6.8 | 119.5 - 125.5 | 6H_480494026 & 6H_481998837 | 15.2 | AWCS799 |
| *Qcrs.caf-2H* | BLUP | IM | 2.8 | 268.5 - 279.5 | 2H_708222741 & 2H_712330343 | 6.8 | AWCS799 |
| *Qcrs.caf-4H* | FCR04 | ICIM | 7.3 | 213.5 - 218.5 | 4H_633922364 & 4H_635066624 | 18.4 | AWCS799 |
|  | FCR04 | IM | 3.0 | 213.5 - 218.5 | 4H_633922364 & 4H_635066624 | 7.0 | AWCS799 |
|  | BLUP | IM | 3.3 | 211.5 - 218.5 | 4H_633922364 & 4H_635066624 | 7.8 | AWCS799 |
| *Qcrs.caf-5H* | FCR02 | ICIM | 3.9 | 95.5 - 97.5 | 5H_264499774 & 5H_300489701 | 16.1 | AWCS799 |
|  | FCR02 | IM | 4.7 | 95.5 - 100.5 | 5H_264499774 & 5H_300489701 | 8.3 | AWCS799 |
| *Qph.caf-6H* | PH | ICIM | 3.7 | 157.5 - 163.5 | 6H_555603298 & 6H_552874297 | 11.7 | Fleet |
| *Qph.caf-7H* | PH | ICIM | 7.4 | 121.5 - 124.5 | 7H_180473897 & 7H_408998941 | 27.8 | AWCS799 |
| *Qhd.caf-5H* | HD | ICIM | 14.4 | 248.5 - 252.5 | 5H_594490721 & 5H_599429072 | 48.9 | AWCS799 |

<sup>a</sup> 'PH' stands for plant height; 'HD' stands for heading date.
<sup>b</sup> 'IM' stands for the analysis conducted using interval mapping; and 'ICIM' stands for the analysis conducted using inclusive composite interval mapping.
<sup>c</sup> Percentage of phenotypic variance explained

The locus on chromosome 2H (*Qcrs.caf-2H*) was identified in the BLUP based IM analysis only. The other two QTL located on chromosomes 4H (*Qcrs.caf-4H*) and 5H (*Qcrs.caf-5H*) were each detected in only one of the four trials. *Qcrs.caf-4H* was mapped in a similar location with a locus reported in an earlier study (Chen et al. 2013a). As none of these loci were consistently detected, they were not further investigated in this study.

Possible effects of *Qcrs.caf-6H* was further assessed in the validating population of Franklin/AWCS799. The marker (6H_497772849) closely linked with *Qcrs.caf-6H* from the mapping population was used to identify individuals with (*RR*) or without (*rr*) the resistant allele in this population. Significant difference was detected for *Qcrs.caf-6H* between *RR* and *rr* group in each of the three trials (Table 4). The average DI value of the lines bearing the resistant allele was 13.9% lower than those with the susceptible allele.

**Table 4.** Disease index of FCR severity of lines possess resistant (*RR*) and susceptible (*rr*) allele of *Qcrs.caf-6H* from the population of Franklin/AWCS799

| Trial <sup>a</sup> | *RR* <sup>b</sup> | *rr* <sup>b</sup> | Difference (%) <sup>c</sup> | *P* value <sup>d</sup> |
| --- | --- | --- | --- | --- |
| FCRV01 | 39.5 | 43.2 | 9.5 | <0.05 |
| FCRV02 | 28.1 | 31.5 | 12.0 | <0.05 |
| FCRV03 | 25.5 | 30.7 | 20.1 | <0.05 |

<sup>a</sup> The three CEF trials conducted were designated as FCRV01, FCRV02 and FCRV03, respectively.
<sup>b</sup> The number of *RR*- and *rr*- genotypes are 50 and 71.
<sup>c</sup> Differences were obtained by comparison between *RR*- and *rr*- genotypes.
<sup>d</sup> '*P* values' were generated with student's *t* test.

### Effect of plant height and heading date on FCR resistance

A QTL controlling HD was identified on chromosome 5H. This QTL explained up to 48.9% of phenotypic variance with a LOD value of 14.4 (Table 3). Two QTL affecting PH were detected and were located on chromosomes 6H and 7H, respectively. The QTL on chromosome 6H could explain up to 11.7% of the phenotypic variance with a LOD value of 3.7 and the other QTL on chromosome 7H could explain up to 27.8% of the phenotypic variance with a LOD value of 7.4 (Table 3). For quantifying possible effects of PH and HD on FCR severity, the BLUP data of the four FCR inoculation trials was analysed against PH and HD data using covariance analysis. The results showed that both PH (Fig. 4) and HD (Fig. 5) have little effect on *Qcrs.caf-6H*.

**Fig. 4.**
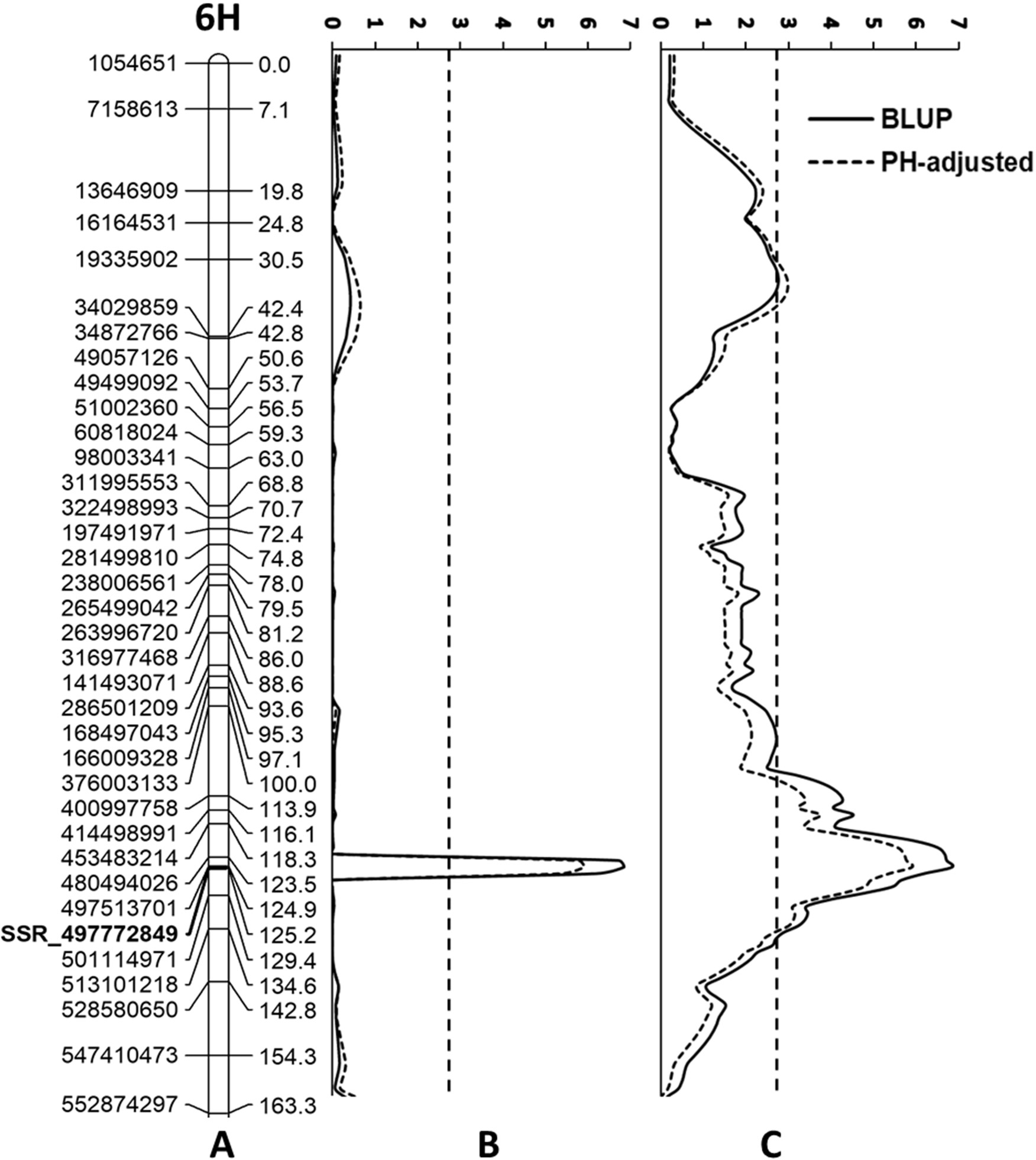
LOD values of *Qcrs.caf-6H* obtained from the BLUP data and post-adjustment by plant height (PH-adjusted). A: Physical position for each marker is shown to the left of the linkage map and distances in centiMorgan (cM) are shown to the right; B: LOD value graph for “ICIM” model; C: LOD value graph for “IM” model. The vertical dotted lines indicate the average significance threshold (LOD=2.7) based on a test of 1,000 permutations. The SSR marker used in validating the QTL is bolded.

**Fig. 5.**
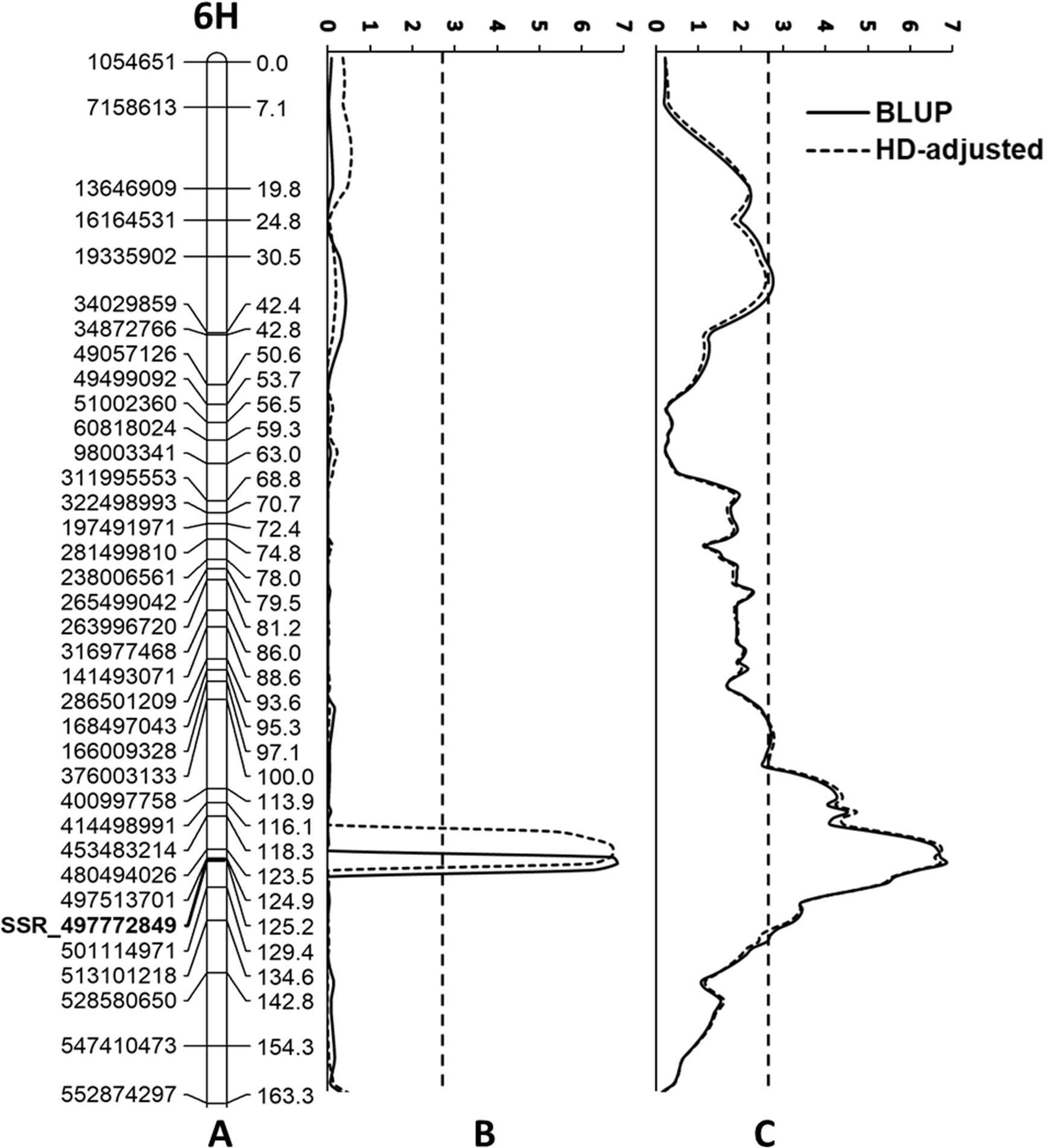
LOD values of *Qcrs.caf-6H* obtained from the BLUP data and post-adjustment by heading date (HD-adjusted). A: Physical position for each marker is shown to the left of the linkage map and distances in centiMorgan (cM) are shown to the right; B: LOD value graph for “ICIM” model; C: LOD value graph for “IM” model. The vertical dotted lines indicate the average significance threshold (LOD=2.7) based on a test of 1,000 permutations. The SSR marker used in validating the QTL is bolded.

## Discussion

In the study reported here, we investigated the genetics of FCR resistance on a barley landrace originating from South Korea. Four loci were detected but only one of them was consistently detected in each of the trials and its effects were also confirmed in a second population. This QTL, designated as *Qcrs.cpi-6H*, located on the long arm of chromosome 6H explaining up to 29.1% of the phenotypic variance. Its effects were detected in both of the populations assessed. This is the first locus conferring FCR resistance identified on this chromosome in barley.

Pyramiding multiple loci into single genetic background has been proved to be effective in improving FCR resistance (Chen et al. 2015; Zheng et al. 2017). The detection of *Qph.caf-6H* further extended the potential of such an approach. However, QTL mapping offers only limited resolution (Paterson et al. 1988) and molecular markers derived from such an approach may not be adequate for effective gene pyramiding. To fulfil the potential of this new QTL, diagnostic markers which can be convenient exploited in breeding programs need to be developed for it. One of the approaches for developing such markers is the use of large populations derived from near isogenic lines. This approach has been successfully exploited in developing co-segregating markers for loci conferring FCR resistance in both wheat (Zheng et al. 2015) and barley (Jiang et al. 2018).

Results from previous studies show that plant height affects FCR severity in both wheat (Collard et al. 2005; Wallwork et al. 2004) and barley (Chen et al. 2013a; Li et al. 2009; Liu et al. 2012a). A histological analysis based on NILs for height also showed that that *F. pseudograminearum* hyphae were detected earlier and proliferated more rapidly during the time-course of FCR development in the tall isolines (Bai and Liu 2015). Some of the earlier studies also showed that FCR severity can be influenced by HD (Liu et al. 2012a). However, the new FCR locus detected in this study does not appear to have strong interactions with either plant height or heading date. The FCR locus does not co-locate with loci controlling either plant height or heading date, and removing the effects of these characteristics from the mapping population has little influence on the magnitudes of the FCR locus detected in any of the trials. It is not clear why this new FCR QTL seems to be different from several of those reported earlier. Nevertheless, the lack of interaction with these characteristics makes the new locus easier to manipulate in breeding programs.

Loci conferring FCR resistance have been reported on 13 of the 21 wheat chromosomes and one of them was located on chromosome arm 6DL (Liu and Ogbonnaya 2015). The locus was detected from a synthetic hexaploid genotype with the use of a single population (Martin et al. 2015). However, none of the markers closely linked with the wheat locus mapped to the corresponding region of the locus detected on the chromosome arm 6HL in this study. This seems to indicate that the two loci are not homoeologous. However, considering the limited resolution of QTL mapping (Paterson et al. 1988), we cannot rule out the possibility that the lack of shared markers could be due to inaccurate locations of either or both of the loci in concern.

As expected, orders for the majority of the markers in the linkage map constructed in this study aligned well with the relative positons of the sequences from which they were generated in the barley genome. It is of interest to note that, without any exception, all the discrepancies involved markers and sequences located in the peri-centromeric region (Fig. S3). It is known that the peri-centromeric regions of the barley chromosomes are characterized by low gene density, low recombination frequencies (Mascher et al. 2017) and high ratios of repetitive sequences (Wicker et al. 2017). Contributions, if any, from these characteristics to the discrepancies are not clear. However, it is not unreasonable to speculate that low recombination frequencies are likely to be less tolerable to incorrect marker scores.

## Supporting information

Fig. S2 A linkage map of barley based on the population of FleetAWCS799

## Author Contributions

C.L., M.Z., Y.W. and Y-L.Z. conceived the research; S.G., Z.Z., H.H., H.S., J.M. and Y.L. conducted experiments and analysed data under the supervision of C.L. and M.Z.; G.S., Z.Z., Y.L. and C.L. wrote the manuscript with contribution from all the authors.

## Acknowledgements

Work reported here was partially funded by the Grains Research and Development Corporation, Australia (Project CFF00010). SG is grateful to China Scholarship Council and University of Tasmania for financial supports during the tenure of his PhD studentship. HH thanks the Henan Institute of Science and Technology and China Scholarship Council for supporting her visit to CSIRO. We are also grateful to Drs Meredith McNeil and Udaykumar Kage (both at CSIRO Agriculture and Food) for their constructive suggestions in preparing the manuscript.

