## Supplementary material for "A novel QTL conferring Fusarium crown rot resistance located on chromosome arm 6HL in barley": TableS1, Fig S1 to S3

Table S1 Chromosome assignment and length of linkage groups, and marker density of the maps based on the population of Fleet/AWCS799

| Chromosome | Number of markers | Length (cM) | Marker density (cM/Marker) |
|---|---|---|---|
| 1H | 84 | 256.3 | 3.1 |
| 2H | 105 | 324.1 | 3.1 |
| 3H | 165 | 356.7 | 2.2 |
| 4H | 82 | 222.0 | 2.7 |
| 5H | 134 | 343.8 | 2.6 |
| 6H | 95 | 199.3 | 2.1 |
| 7H | 76 | 262.5 | 3.5 |
| Total | 741 | 1964.7 | 2.7 |

Fig. S1 Distribution of disease index values for the four FCR trials and the BLUP data from the mapping population Fleet/AWCS799.

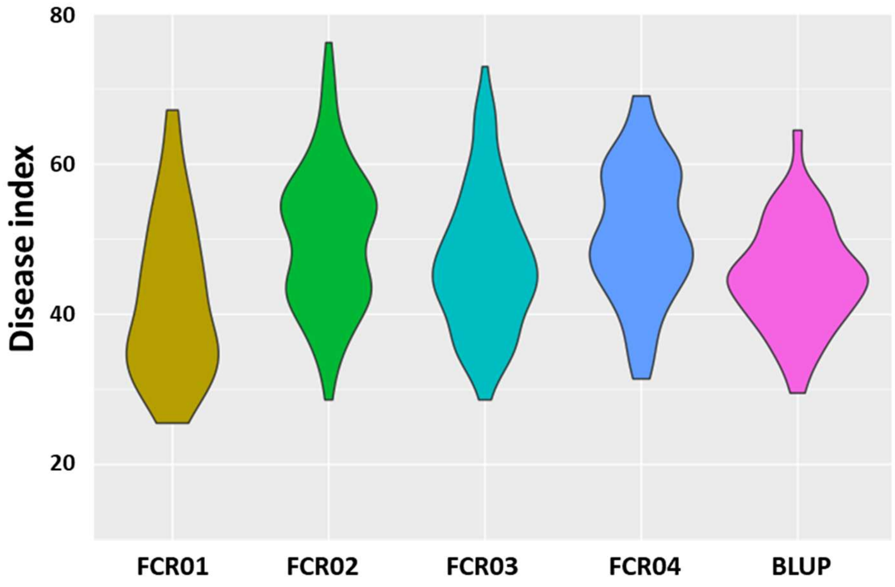

Fig. S2 A linkage map of barley based on the population of Fleet/AWCS799 (See Additional file)

Fig. S3 Syntenic relationships for the mapped markers between the genetic and physical positions. Horizontal-axis represents genetic distance (cM) and vertical-axis represent physical distances (base pairs). Peri-centromeric regions were highlighted in gray shadow.

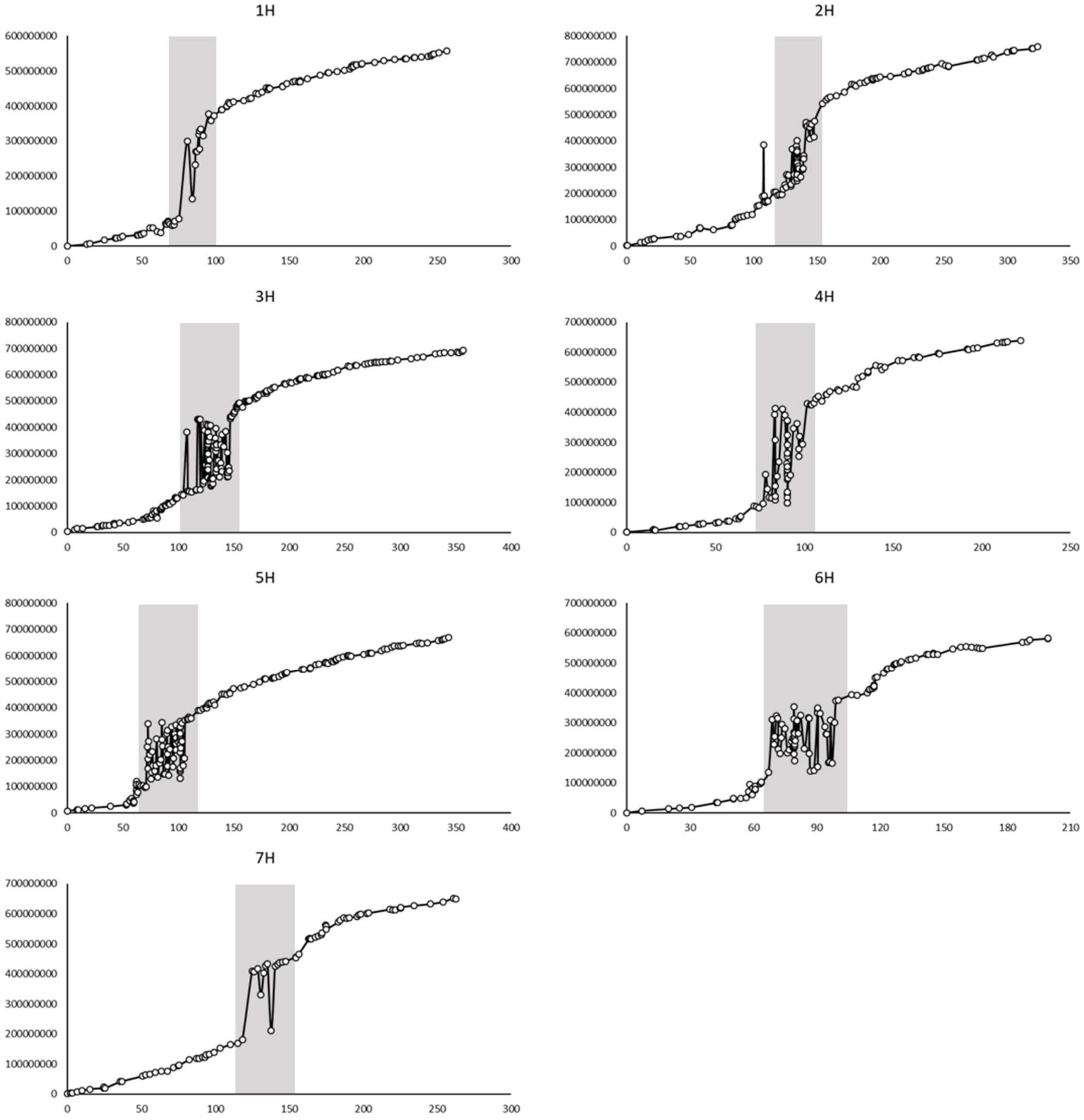
