## Supplementary figures and images for "A novel QTL conferring Fusarium crown rot resistance located on chromosome arm 6HL in barley"

### Fig. S2 A linkage map of barley based on the population of FleetAWCS799.jpg

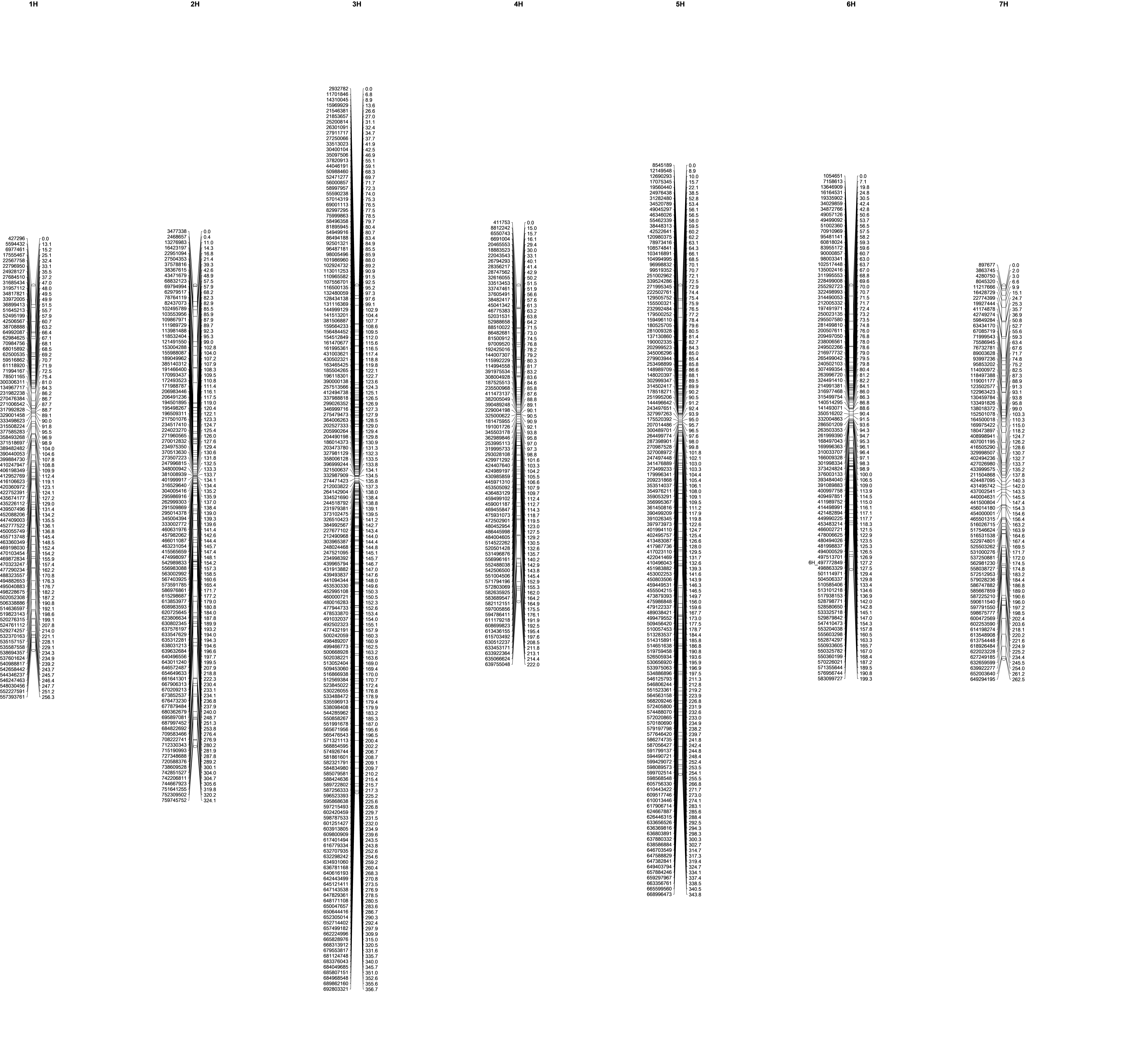
